# Homogeneous network inference for subtype-stratified master regulator analysis in cancer

**DOI:** 10.64898/2026.11.07.765099

**Authors:** Hugo Tovar, Camila Barrios-Cervantes, Daniel Pérez-Calixto, Enrique Hernández-Lemus

## Abstract

Master-regulator analysis of 34 TCGA tumour cohorts (cancer × molecular subtype; *n* between 27 and 562 tumours) requires one regulatory network per cohort. Mutual-information (MI) networks inferred with ARACNe3 are the usual choice, but they are not available for the smallest cohorts; filling the gap with correlation networks would mix two methods inside the very comparison between subtypes that the analysis is meant to make. We therefore infer every network with a single procedure, *fixed-size Spearman regulons* (FSR): rank correlation between regulator and target, a fixed effect-size threshold |*ρ*_*S*_| ≥ 0.2, the per-target data-processing-inequality (DPI) pruning implemented in corto, bootstrap support, and truncation to the 100 best-supported targets per regulator. We quantify what this choice concedes and what it provides, labelling each claim as a standard result, a heuristic, or an empirical result. Rank correlation is more powerful than the MI estimators we simulated for *n ≲* 200 but cannot see non-monotone dependence, which in a synthetic worst case costs at most about 0.05–0.10 of the master regulators recovered; the per-target DPI is a heuristic with a measurable disagreement rate with the MI-based one; and pairwise dependence is not direct interaction, for MI and correlation alike. Where both methods are feasible (*n* ≥ 95) they overlap in 0.7 of the top-20 regulators, the same as two filterings of the same ARACNe3 network, although the rank correlation of the full score vector shows a measurable method effect in the tail. Subsampling large real cohorts indicates that networks inferred from fewer than 60 tumours do not reliably reproduce their own cohort’s master regulators, that 60–100 is a grey zone, and that a cancer-wide network does not rescue small subtypes. We present FSR as an adequate and uniform choice for multi-cohort analysis, not as a statistically superior one.

## 1 Introduction

Master regulator analysis (MRA) is a systems-biology approach for identifying the transcription factors (TFs) that control the transcriptional state of a tumour or a tumour subtype [19, 20]. Instead of testing genes one at a time, MRA first infers a regulatory network, with each TF linked to the targets whose expression it is inferred to influence, and then asks whether a TF’s targets change coherently in a phenotype-specific signature, such as tumour versus normal tissue. A TF whose regulon is significantly enriched in the signature is a candidate master regulator. Because a regulator is scored through many targets at once, the approach is more robust to noise in individual genes than differential expression alone, and it can recover regulators whose own mRNA levels barely change, as their activity is captured through their targets [21]. This is why MRA has been used to find drivers of tumour initiation and progression, to compare the regulatory architecture of cancers, and to prioritise candidate therapeutic vulnerabilities [22]. Its results depend on the regulatory network behind them, so how the network is inferred is a methodological choice that deserves scrutiny.

That choice becomes critical in multi-cohort analyses. Stratifying TCGA by cancer and molecular subtype yields cohorts whose sizes differ by more than an order of magnitude (here *n* from 27 to 562). The standard mutual-information network method, ARACNe3, is built on MI estimated by adaptive partitioning and is designed for the larger end of that range; for the smaller cohorts the practical alternative is a correlation network. Two designs are then possible:

1. *Heterogeneous:* MI networks where the cohort is large enough, correlation networks else-where.
2. *Homogeneous:* one procedure for every cohort.

The question is not which dependence measure is statistically best in the abstract, but which design supports the comparisons the analysis is built for, namely differences in master regulators between subtypes of one cancer and between cancers, so that those differences reflect biology and not the algorithm that happened to be feasible for each cohort. We show in Section 3 that changing the network method alters the top of the master-regulator ranking by an amount of the same order as changing the subtype, so that a heterogeneous design confounds the comparison of interest. We therefore adopt a homogeneous design built on rank correlation, fixed-size Spearman regulons (FSR), which is the only option of the two that produces a network at every *n*.

The contribution of this work is a quantified account of the trade-off, not a claim of superiority. We examine (i) the statistical efficiency of rank correlation relative to MI estimators, (ii) its blindness to non-monotone dependence and the consequence for the ranking, (iii) the validity of per-target DPI pruning on correlations, (iv) the gap between pairwise dependence and direct interaction, which MI shares, (v) the agreement between FSR and ARACNe3 where both are feasible, and (vi) a minimum cohort size for a subtype-specific network. Each claim is labelled as a *standard result* (proved elsewhere), a *heuristic*, or an *empirical result* (simulation or real data), and Section 8 collects the labels.

## 2 Setting and procedures

Let *X ∈* ℝ^*p×n*^ be the expression matrix (log_2_(TPM+1)) of a cohort with *p* ≈ 14 000 genes and *n* tumours, and let *T* be the set of transcription factors (Lambert et al. [9]; 1,204 are present in the COAD CMS2 matrix used for the examples). For each *t ∈ T* we need a *regulon R*_*t*_ = {(*g, w*_*tg*_, *m*_*tg*_)}: targets *g* with a weight *w*_*tg*_ *∈* (0, 1] and a mode *m*_*tg*_ *∈* [−1, 1]. Given a signature *s ∈* ℝ^*p*^ (the DESeq2 Wald statistic, tumour vs. normal; Love et al. [12]), NaRnEA (Griffin et al. [6]) computes a normalised enrichment score NES_*t*_ that combines a directed term (linear in *w*_*tg*_ *m*_*tg*_) and an undirected term (in *w*_*tg*_).

We compare two procedures for building *R*_*t*_:

- **ARACNe3**: *I*(*t*; *g*) estimated by adaptive partitioning on subsamples (63 % without replacement, 100 subnetworks), thresholded by FDR against a null of shuffled pairs, pruned by the DPI over regulator–regulator–target triangles, and consolidated by support across subnetworks; *m*_*tg*_ is the Spearman correlation.
- `corto` (Mercatelli et al. [16]): Pearson correlation *ρ*_*P*_ (*t, g*), here computed on ranks and therefore equal to *ρ*_*S*_; a threshold |*ρ*| ≥ *r*_min_; and DPI pruning in a per-target form (`corto` 1.4.0, `dpi = “target”`): for each target *g* and each bootstrap, among the regulators that pass the threshold only arg max_*t*_ |*ρ*(*t, g*) | is retained (Section 5.1 relates this rule to the triangle DPI). *w*_*tg*_ is the fraction of bootstraps, including the unresampled data, in which (*t, g*) is retained; *m*_*tg*_ = *ρ*.

Both networks are truncated to the 100 best-supported targets per regulator. We call the `corto`-based procedure *fixed-size Spearman regulons* (FSR) and the truncated ARACNe3 network *ARACNe3-top100*. After truncation the two procedures differ only in the dependence measure and in the form of the DPI pruning.

### Evaluation

Three kinds of evidence are used: simulations of dependence measures and chains (Sections 4.1–5.1), synthetic networks with known master regulators and a synthetic block of correlated TFs (Sections 4.2, 5.2 and 6; the generators are described in the Supplementary Note), and real TCGA cohorts (Sections 3–7). Agreement between two rankings of master regulators is the overlap of their top 20 regulators by |NES| (Section 7 checks *k ∈* { 5, …, 50 }). Agreement is a measure of consistency, not of correctness.

## 3 Why a single procedure

The choice between a homogeneous and a heterogeneous design depends on how large the effect of the method is compared with the effect of the biology the analysis is designed to detect. Three yardsticks, all measured as the top-20 overlap, give the scale (real cohorts; details and figures in the Supplementary Note):

- *Null*. The cohort’s own signature scored with the network of another cancer shares 0.35 of the top 20. The overlap expected by chance is negligible (0.0164 at *k* = 20), so a sizeable part of the ranking is independent of the network, being driven by the signature and by the set of TFs.
- *Subtype effect*. Between two subtypes of the same cancer, the overlap is 0.55 when only the network changes, 0.6 when only the signature changes, and 0.45 when both change. The turnover caused by changing the network between subtypes is 0.45.
- *Method effect*. With the same signature and the same cohort, FSR and ARACNe3-top100 differ by a turnover of 0.3, about two thirds of the turnover caused by changing the network between subtypes and about half of the total subtype effect (0.55). The rank correlation of the full NES vector is 0.926 between methods and 0.985 between two filterings of the same ARACNe3 network.

The method effect is smaller than the subtype effect but of the same order and not negligible. Mixing methods across subtypes would therefore confound the comparison the analysis is designed to make: part of the apparent difference between a large and a small subtype of the same cancer would be a difference between algorithms. This is the argument for a homogeneous design; the remainder of the article asks what rank correlation concedes and whether it can be used at every *n*.

## 4 The dependence measure

### 4.1 Statistical efficiency of rank correlation

#### Standard results

Under bivariate normality, the Pitman asymptotic relative efficiency of *r*_*S*_ with respect to *r*_*P*_ *for testing ρ* = 0 is (3*/π*)^2^ = 0.912; the loss is about 9 % in efficiency for that test. For *estimation, r*_*S*_ does not estimate *ρ* but 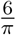 arcsin(*ρ/*2), so a comparison of mean squared errors with respect to *ρ* conflates bias and variance; the comparable quantities are the variance and the power. Under the same model, 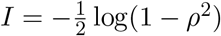 is a monotone function of |*ρ*| : under normality MI does not rank pairs better than correlation and only adds estimation variance. This no longer holds away from normality; non-monotone dependence is treated in Section 4.2.

#### Simulation (Fig. 1)

Pairs (*x, y*) with *ρ* = 0.4, 200 replicates for each of *n* = 27, 50, 100, 200 and 400, under three noise models calibrated against the tails of real expression data. The per-gene excess kurtosis in COAD CMS2 (log_2_TPM) has median 0.281, 90th percentile 1.98 and 99th percentile 5.62, with a fraction 0.0396 of genes above 3 (panel C). The models are: Gaussian; Student *t* with 5 degrees of freedom (excess kurtosis 6, above the observed 99th percentile; a *t*_3_ would be unrealistically heavy-tailed); and Gaussian with 3 % outliers, the dominant pattern in RNA-seq data. We record the standard deviation of *r*_*P*_ and *r*_*S*_, and the power to detect dependence at the 5 % level against a permutation null, for Pearson, Spearman and three MI estimators: the naive 8-bin estimator, Miller–Madow [17] with 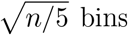 bins, and the *k*-nearest-neighbour estimator of Kraskov, Stögbauer and Grassberger (KSG) [7] with *k* = 3.

**Figure 1.**
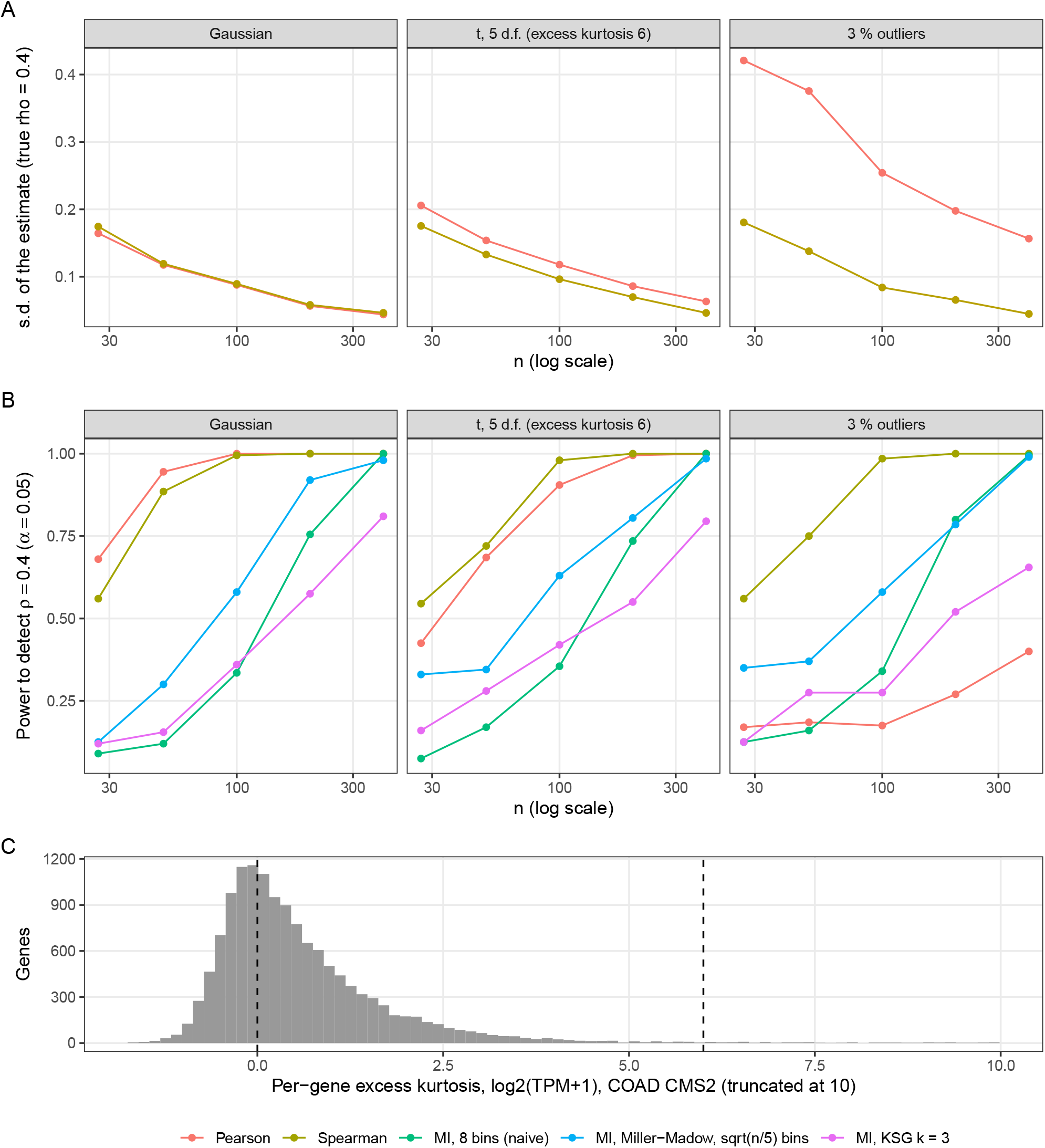
(A) Standard deviation of *r*_*P*_ and *r*_*S*_ with *ρ* = 0.4. (B) Power at the 5 % level for each measure. (C) Per-gene excess kurtosis in COAD CMS2; vertical lines at 0 (Gaussian) and 6 (*t*_5_).

**Figure 2.**
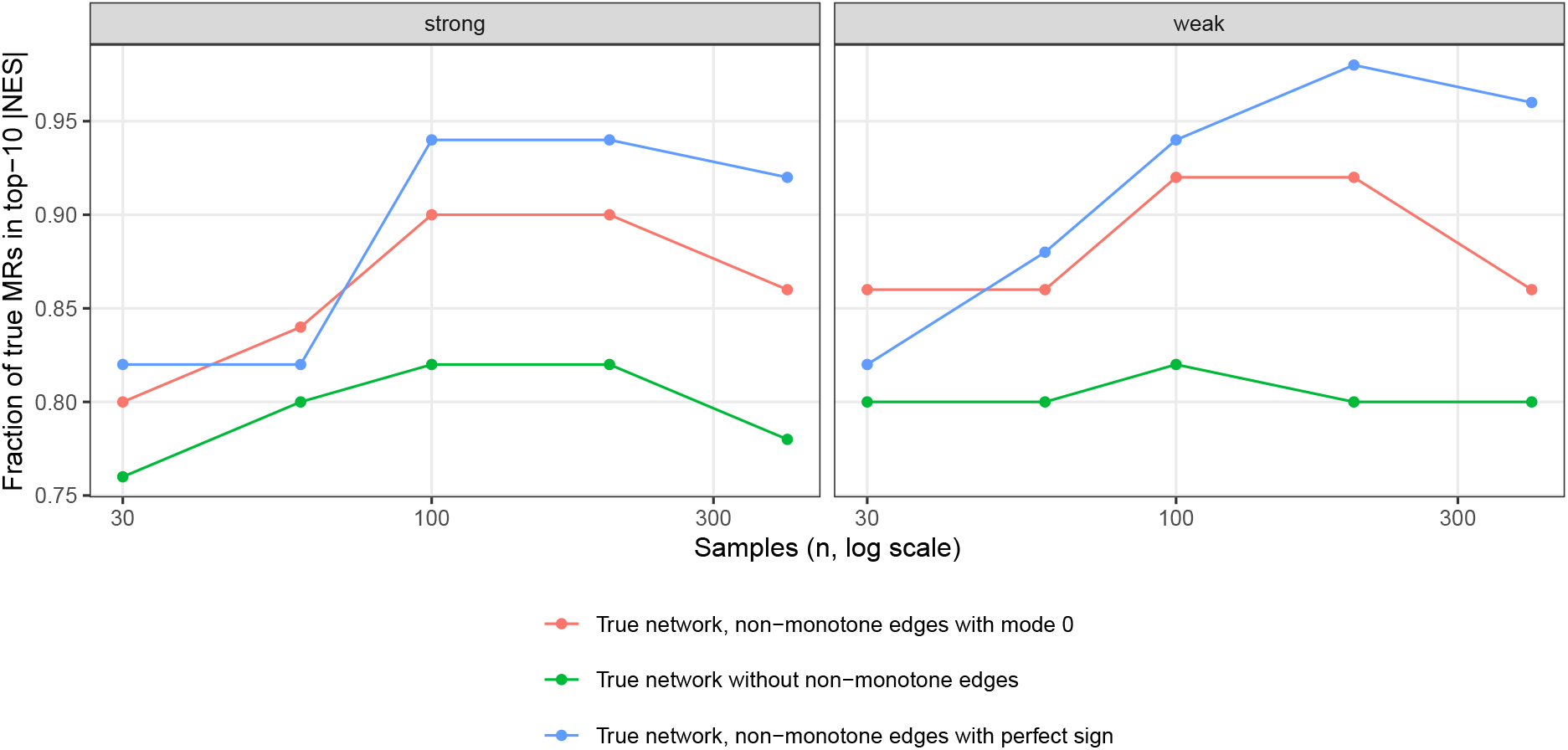
Recovery of true master regulators with the true synthetic network, depending on how its non-monotone edges are represented.

- Gaussian: variance ratio Var(*r*_*P*_ )*/*Var(*r*_*S*_) = 0.934 (mean over *n*; this is not the Pitman efficiency, which is defined as *ρ* → 0). Power at *n* = 100: Pearson 1, Spearman 0.995, MI-KSG 0.36, MI Miller–Madow 0.58, naive MI 0.335.
- *t*_5_: standard deviation at *n* = 100 of 0.118 (Pearson) versus 0.0963 (Spearman); power 0.905 versus 0.98.
- Outliers: standard deviation at *n* = 400 of 0.156 (Pearson) versus 0.0448 (Spearman); power 0.4 versus 1.
- MI: with a reasonable estimator (KSG or Miller–Madow) the power at *n* = 100 is 0.36–0.58 under normality, and 0.575–0.92 at *n* = 200: well below Spearman (0.995 and 1) but far from zero. The naive 8-bin estimator at *n* = 27 (64 cells) has a Miller–Madow bias of ≈ (*B* − 1)^2^*/*2*n* ≈ 0.9 nats; it is shown only to illustrate the size of the estimation problem.

*Remark* 1. None of these is the ARACNe3 estimator (adaptive partitioning), and the simulation uses a single effect size, *ρ* = 0.4, whereas the edges of real networks span a wider range (Fig. 3). The direct comparison with ARACNe3 is the empirical one in Section 6. This section establishes that, for *n ≲* 200, the MI estimators simulated lose power relative to Spearman, and that Spearman is more efficient than Pearson as soon as the data have heavy tails or outliers.

**Figure 3.**
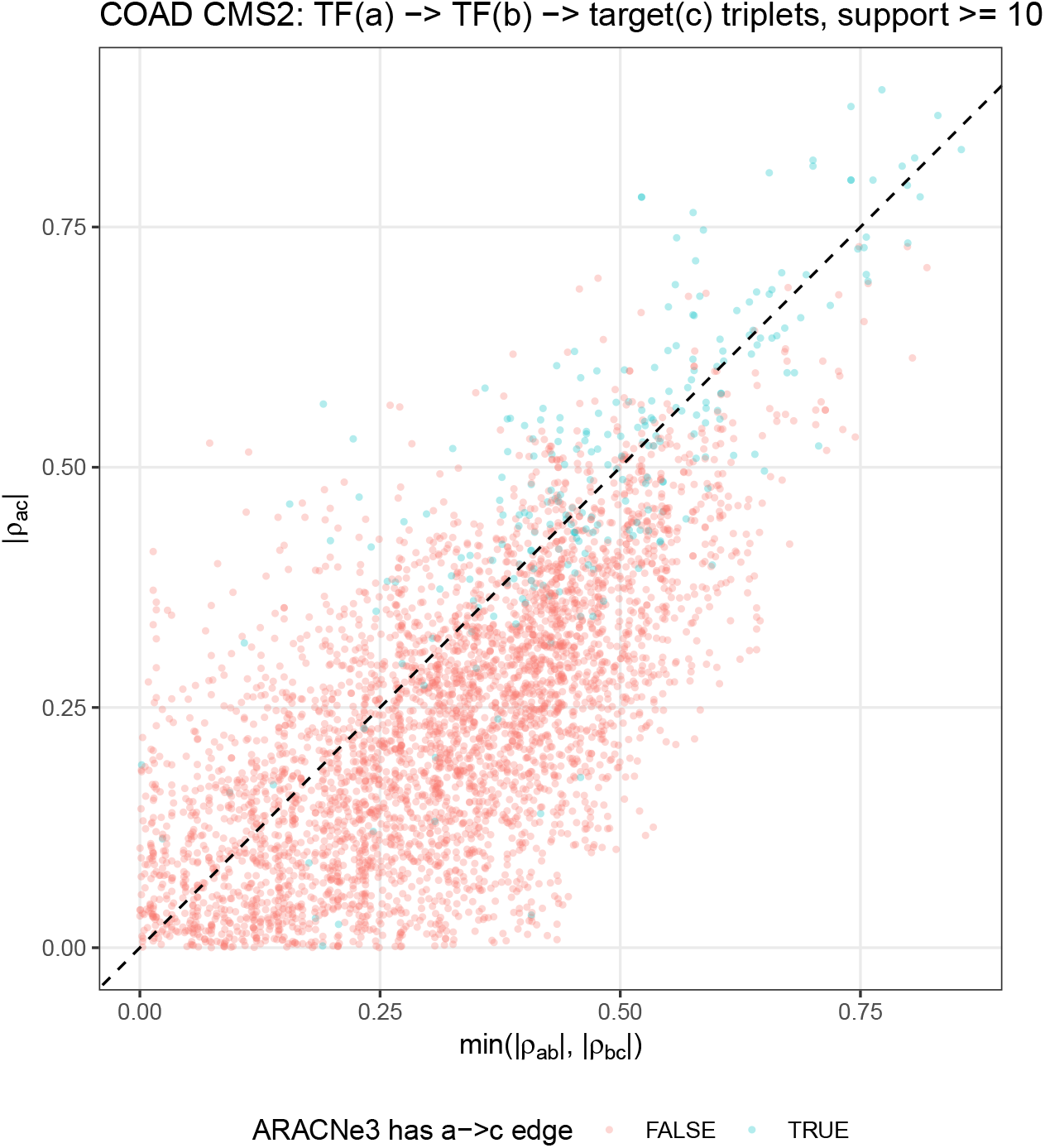
TF→TF→target triplets in the ARACNe3 network of COAD CMS2. Points above the diagonal violate |*ρ*_*ac*_| ≤ min(|*ρ*_*ab*_|, |*ρ*_*bc*_|). Colour: whether ARACNe3 also infers the direct edge *a* → *c*.

### 4.2 Monotone and non-monotone dependence

By construction, *ρ*_*S*_(*x, f* (*x*)) = ± 1 for every monotone *f* and ≈ 0 for an even (U-shaped) *f* . MI is invariant under any bijective reparametrisation and is zero if and only if the variables are independent (Cover and Thomas [2]). Two questions follow: whether non-monotone regulatory dependencies are present in tumour data, and what they cost the master-regulator ranking if FSR cannot see them.

#### Presence

In synthetic data in which 20 % of the true edges are non-monotone, ARACNe3 recovers them at *n* = 400 (recall 0.82 under strong signal) and FSR does not (0.0733). In the real ARACNe3 networks their presence can be neither established nor excluded: the fraction of edges with |*ρ*_*S*_| *<* 0.2 among those with support ≥ 10 is 0.043 in the real tumours and 0.0582 in the synthetic positive control with 20 % non-monotone edges (*n* = 400; 0.000333 at *n* = 100). The statistic is nearly blind to such edges over the range of *n* of our cohorts, and the question remains open. The simulated non-monotone edges are the worst case for correlation, a symmetric U with *ρ*_*S*_ ≈ 0; real non-monotone responses are usually asymmetric and retain |*ρ*| *>* 0, which favours correlation-based inference more than this analysis shows.

#### Effect on the ranking (Fig. 2)

Because the directed term of the NES is linear in *m*_*tg*_, removing edges with |*m*_*tg*_| ≈ 0 from an *inferred* regulon can change little by construction, so such a test would be tautological. Instead we use the *true* synthetic network, free of inference error, in three versions: with the non-monotone edges removed; with them present and given mode 0; and with them assigned a “perfect” sign |*m*| = 1. The three versions map onto the methods as follows. FSR does not detect the edges (recall above), so what it delivers is closest to the first version. An MI method detects them but, because its mode is the Spearman correlation, assigns them mode ≈ 0: the second version. The third is an oracle, information that no correlation can provide and that bounds what a scoring method able to exploit non-monotone dependence could gain. Fraction of true master regulators in the top 10, mean of 5 replicates:

|  | without non-monotone (FSR-like) | with mode 0 (MI-like) | with perfect sign (oracle) |
|---|---|---|---|
| strong, *n* = 400 | 0.78 | 0.86 | 0.92 |
| weak, *n* = 400 | 0.8 | 0.86 | 0.96 |
| strong, *n* = 100 | 0.82 | 0.9 | 0.94 |

Non-monotone edges do contribute to the score. The gap between the first two columns, roughly 0.05–0.10 of the recovered fraction, is therefore the most that FSR can lose to an MI method through this mechanism in this worst case (20 % of edges, all symmetric U-shaped, true network). The real loss depends on a prevalence that we cannot estimate. The gap between the second and third columns is one to two standard errors (standard error of the fraction: 0.02–0.0775, with 5 replicates), so it is not resolved by this experiment; it indicates that an MI method scored through the current signed statistic captures most of what a signed representation would add, and that the rest is not reachable by changing the dependence measure alone. This is a property of NaRnEA, not a general argument; a change to an unsigned enrichment method would require revisiting the choice.

## 5 Pruning and the limits of pairwise networks

### 5.1 The data processing inequality on correlations

For a Markov chain *X* →*Y* → *Z, I*(*X*; *Z*) ≤ min{*I*(*X*; *Y* ), *I*(*Y* ; *Z*) } (Cover and Thomas [2]). ARACNe uses this inequality to remove, in every triangle of edges that pass the threshold, the edge with the smallest MI. `corto` also applies DPI pruning, in a simplified per-target form. With this rule (`dpi = “target”`, the default in `corto` 1.4.0) every target *c* keeps, in each bootstrap, only the regulator with the largest |*ρ*(·, *c*) | among those that pass the threshold. For two candidate regulators *a* and *b* of *c*, this is the triangle DPI under the assumption that the regulator–regulator edge *a*–*b* is never the weakest of the triangle: `corto` never computes *ρ*(*a, b*). Two consequences follow. In a chain *a* → *b* → *c*, the rule removes *a* → *c* exactly when |*ρ*_*ac*_| *<* |*ρ*_*bc*_|, which is the half of the inequality that identifies the indirect edge. When *a* and *b* are independent co-regulators of *c*, no triangle exists and ARACNe keeps both edges, whereas `corto` keeps only the stronger one in each bootstrap; the weaker co-regulator survives only through bootstrap variability, which is what its weight *w*_*tg*_ measures. The rule is therefore more aggressive than the triangle DPI, and its weight measures how often an edge wins this competition, not the stability of the edge itself.

Whether the rule removes the indirect edge depends on whether |*ρ*_*bc*_| *>* |*ρ*_*ac*_| in a chain, that is, whether the direct regulator outscores the indirect one.

**Proposition 1**. *If Y* = *αX* + *ε*_1_ *and Z* = *βY* + *ε*_2_, *with ε*_1_, *ε*_2_ *independent of each other and of X and with finite variances (linearity and independent noise, with no distributional assumption), then ρ*_*XZ*_ = *ρ*_*XY*_ *ρ*_*Y Z*_. *In particular* |*ρ*_*XZ*_| ≤ min(|*ρ*_*XY*_ |, |*ρ*_*Y Z*_|); *moreover* |*ρ*_*XZ*_| = |*ρ*_*Y Z*_| *only if* |*ρ*_*XY*_ | = 1 *or ρ*_*Y Z*_ = 0.

*Proof*. Cov(*X, Z*) = *αβ* Var(*X*); normalising, 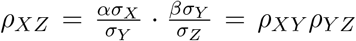 Since |*ρ*_*XY*_ | ≤ 1 and |*ρ*_*YZ*_| ≤ 1, |*ρ*_*XZ*_| ≤ |*ρ*_*YZ*_| and |*ρ*_*XZ*_| ≤ |*ρ*_*XY*_ |, and the equality |*ρ*_*XZ*_| = |*ρ*_*YZ*_| requires |*ρ*_*YZ*_| (1 − |*ρ*_*XY*_ |) = 0.

*Remark* 2. The proposition concerns *ρ*_*P*_ and does not require normality. Carrying it over to *ρ*_*S*_, which is what FSR compares, requires *ρ*_*S*_ to be a monotone function of *ρ*_*P*_ ; under bivariate normality 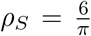 arcsin(*ρ*_*P*_ */*2) and the inequality is preserved. Normality enters at this step only.

Outside this regime the rule can fail. We quantify it with two statistics: the triangle inequality

|*ρ*_*ac*_| ≤ min(|*ρ*_*ab*_|, |*ρ*_*bc*_|), for comparison with the ARACNe DPI, and the comparison that the per-target rule actually makes, *P* (|*ρ*_*ac*_| *>* |*ρ*_*bc*_|).

- **Simulated chains** (*n* = 400, 200 replicates) with *f ∈* { linear, sigmoid, quadratic, absolute value} at each link. With both links monotone, the fraction of violations of the triangle inequality is 0 (linear–linear), 0 (sigmoid–sigmoid) and 0 (linear–sigmoid). With one non-monotone link it is of the order of one half (0.425 quadratic–linear; 0.425 abs–abs). This value is not a failure rate of the per-target rule: with a non-monotone link, *ρ*_*S*_ does not measure the dependence along that link, and the statistic then compares two correlations that are both near zero.
- **Real triplets** (Fig. 3): 4,186 triplets *a* → *b* → *c* (up to 5 targets per pair *a, b*; 838 distinct pairs, so the triplets are not independent) taken from the ARACNe3 network of COAD CMS2 with support ≥10. The triangle inequality is violated in 0.255 of them: 0.236 when ARACNe3 does not infer the direct edge *a* → *c* and 0.558 when it does. The per-target rule depends on |*ρ*_*ac*_| *>* |*ρ*_*bc*_|, which occurs in 0.199 of all triplets, 0.182 of those without a direct edge and 0.471 of those with one.

*Remark* 3. In the triplets that MI itself treats as chains (no direct edge), the indirect regulator outscores the direct one in a fraction 0.182 of cases. This is a *disagreement rate* with the MI-based DPI, not an error rate against a known truth, because the real triplets have no ground truth; it shows that the per-target rule is a heuristic whose decisions differ measurably from those of the triangle DPI. The violation rate of 0.558 for triplets with a direct edge partly reflects selection, since ARACNe3 retained *a* → *c* precisely when *I*(*a, c*) was not the smallest edge of the triangle. These triplets are not pure chains either: common confounders such as tumour purity or proliferation create *ρ*_*ac*_ without a chain, and the MI-based DPI is equally blind to them.

#### The triangle rule

`corto` ≥ 1.4.0 also offers the triangle form of the DPI (`dpi = “triangle”`), closer to ARACNe: the edge *a*–*c* is removed only if another regulator *b* has both |*ρ*(*b, c*) | and |*ρ*(*a, b*) | larger than |*ρ*(*a, c*) | . We evaluated it as an alternative to the per-target rule (Supplementary Note, Section 3.9). It retains 2.82–4 times as many edges before truncation and leaves fewer indirect edges in small samples (-6.1 and -13.9 indirect false positives per 100 edges at *n* = 60 and *n* = 100), but it did not meet the criterion fixed in advance, an increase of 0.05 in the fraction of true master regulators in the top 10 (mean difference -0.008, range -0.06 to 0.06 across regimes and sample sizes), it lost edge precision at *n* = 400 (-0.527 with strong and -0.211 with weak signal), and it took 35.5–48.7 min per network against 6.16–8.52 min for the per-target rule. In 5 real cohorts the two rules share 0.6–0.9 of the top-20 master regulators; the triangle rule agrees somewhat more with ARACNe3-top100 (0.7–0.85 against 0.5–0.75), which is consistency with another method, not evidence of correctness. We therefore keep the per-target rule.

### 5.2 Pairwise dependence and direct interaction

#### Standard result

Pairwise dependence is not direct interaction. For a Gaussian vector with precision matrix Θ = Σ^*−*1^, the partial correlation of *i* and *j* given all other variables is 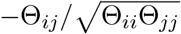, and *X*_*i*_ *⊥ X*_*j*_ |*X*_rest_ if and only if Θ_*ij*_ = 0 (Lauritzen [10]), whereas the marginal correlation Σ_*ij*_ is in general non-zero for any two variables connected by a path in the graph of Θ. Recovering couplings from correlations requires inverting the correlation matrix (partial correlation, or the inverse Ising problem; Schneidman et al. [18]). Thresholding pairwise correlations yields a *co-expression* network, not a regulatory one, and DPI pruning is a heuristic to move it towards the latter. The same holds for ARACNe: pairwise MI has the same limitation and its triangle DPI is likewise a heuristic. The empirical questions are how much each heuristic removes and what the conditional alternative (regularised partial correlation, the graphical lasso; Friedman et al. [5]) achieves in the *p ≫ n* regime.

#### Simulation (Fig. 4)

A synthetic block of 40 TFs (two modules of mutually correlated regulators, latent *ρ* = 0.5) and 600 targets, *p* = 640, with linear Gaussian dependence. For every 100 selected edges we count the *indirect* false positives (the TF shares a module with a true regulator of the target). Since the number of selected edges affects precision and the indirect count, all methods are compared *at the edge count of FSR*, ARACNe3 included (ranked by support, then by MI). At *n* = 100 ARACNe3 produces fewer edges than FSR in the block, so its edge count cannot be matched: the “ARACNe3 by support” row and the “ARACNe3-top100” row coincide there.

**Figure 4.**
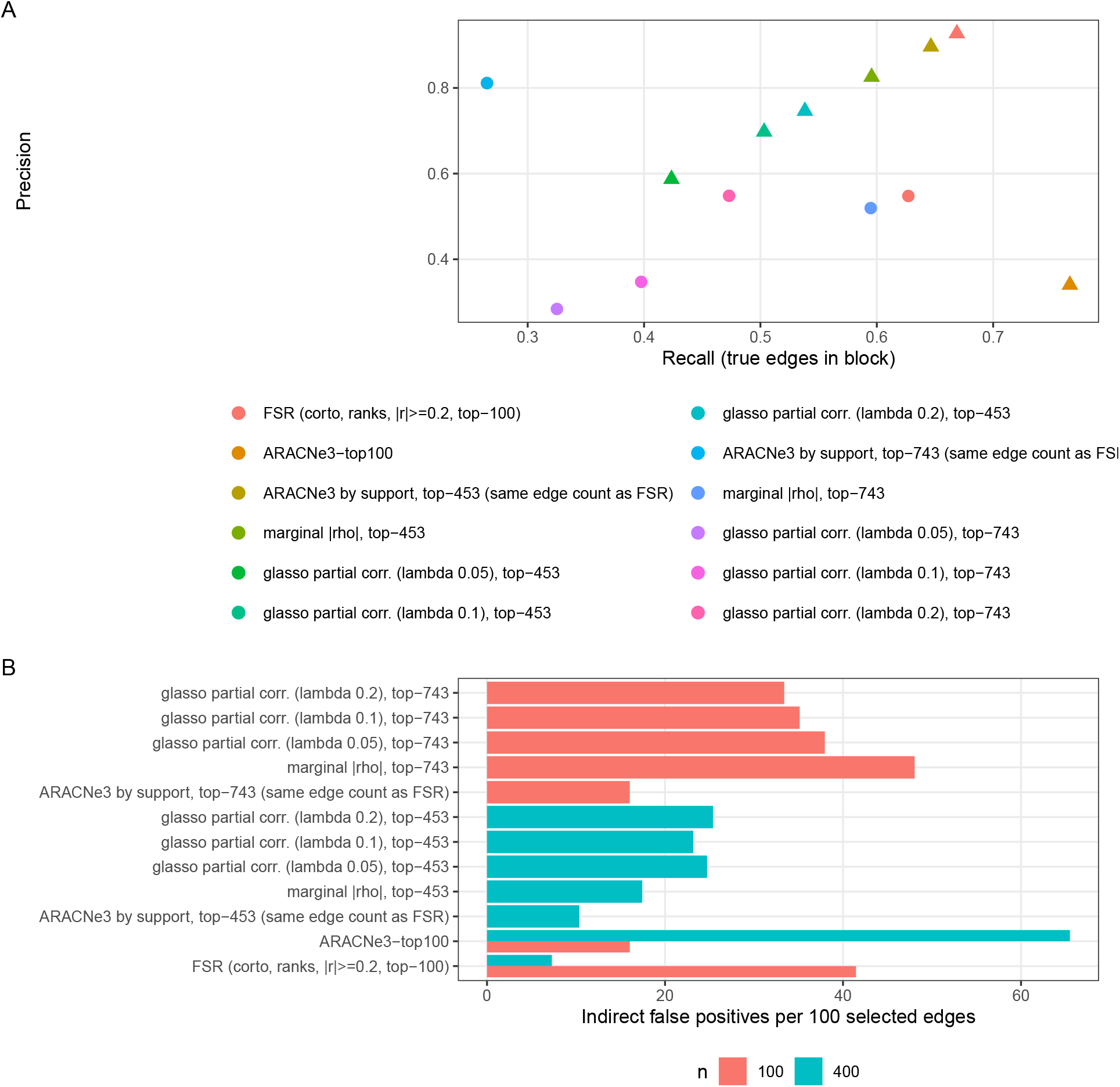
Indirect edges in a synthetic block of correlated TFs. (A) Precision–recall; (B) indirect false positives per 100 selected edges. Except for ARACNe3-top100 with all its edges, every method selects as many edges as FSR where it can reach that count (ARACNe3 at *n* = 100 and the graphical lasso with *λ* = 0.2 at *n* = 100 select fewer).

|  | *n* = 400 |  |  | *n* = 100 |  |  |
|---|---|---|---|---|---|---|
| Method (same edge count as FSR, where possible) | prec. | recall | indir./100 | prec. | recall | indir./100 |
| FSR (corto, ranks, top-100) | 0.927 | 0.669 | 7.28 | 0.548 | 0.627 | 41.4 |
| ARACNe3 by support | 0.896 | 0.646 | 10.4 | 0.811 | 0.265 | 16 |
| Marginal correlation \|*ρ*\| | 0.826 | 0.596 | 17.4 | 0.52 | 0.595 | 48 |
| Partial correlation (glasso, *λ* = 0.1) | 0.698 | 0.503 | 23.2 | 0.347 | 0.398 | 35.1 |
| ARACNe3-top100 (*all* its edges) | 0.34 | 0.766 | 65.5 | 0.811 | 0.265 | 16 |

Four observations:

1. *At n* = 400, *equivalent ordering of edges*. At equal edge counts, the per-target DPI of `corto` and the support ranking of ARACNe3 remove indirect edges at comparable rates (7.28 and 10.4 per 100 selected edges): they order the edges in a similar way. They differ in how many edges they retain. The per-target rule leaves a recall of 0.669 in the block, whereas ARACNe3-top100, at the cap of 100 targets per regulator, retains many more edges (recall 0.766) with far more noise (precision 0.34, 65.5 indirect per 100). Any apparent advantage of either method in indirect edges at unequal edge counts is an artefact of network size.
2. *At n* = 100, *the per-target rule is not equivalent to ARACNe3*. ARACNe3 is far more conservative: higher precision (0.811 vs. 0.548), fewer indirect edges (16 vs. 41.4 per 100), and much lower recall (0.265 vs. 0.627). FSR also has more indirect edges than the graphical lasso row (35.1), although with higher precision and recall. This is the usual precision– recall trade-off. For NaRnEA, which aggregates 100 weighted targets, recall matters more than precision (Supplementary Note, Section 3.4).
3. *Against marginal correlation and the graphical lasso*. FSR has higher precision and fewer indirect edges than marginal correlation |*ρ*| at both *n*: by a wide margin at *n* = 400 (precision 0.927 vs. 0.826; 7.28 vs. 17.4 indirect per 100), and by a small one at *n* = 100 (0.548 vs. 0.52; 41.4 vs. 48). FSR has higher precision than the graphical lasso for every *λ* at *n* = 400 and for *λ* ≤ 0.1 at *n* = 100; with *λ* = 0.2 at *n* = 100 the precision is the same (0.548), with fewer edges selected and lower recall. FSR has fewer indirect edges than the graphical lasso only at *n* = 400. The graphical lasso is at a stated disadvantage here: *p* = 640 *> n, λ* is fixed rather than selected (by StARS [11] or EBIC [4]), and the targets in the block have regulators outside it that the method does not observe. It is the version of the conditional approach that fits the *p ≫ n* regime without further assumptions, not its best version. With the 14 000 real genes the obstacle is computational cost and conditioning, not principle (nodewise regression in the manner of Meinshausen and Bühlmann [15] would scale); we have not pursued it.
4. *Regime*. The block is linear and Gaussian, the regime most favourable to correlation-based methods; it supports statements about indirect edges in that regime only.

## 6 Agreement with ARACNe3 where both are feasible

The preceding sections concern properties of the networks. The output of the pipeline is a ranking of master regulators, for which two further measurements are available (details and figures in the Supplementary Note):

- *Real cohorts*. In the 15 cohorts with *n* ≥ 95, FSR and ARACNe3-top100 have a median top-20 overlap of 0.7, the same as between two filterings of the same ARACNe3 network (0.7). The two procedures agree on most, not all, of the head of the ranking. The rank correlation of the full NES vector is 0.926 between methods and 0.985 within ARACNe3: there is a measurable method effect in the tail of the ranking, not at its head.
- *Synthetic data* with known master regulators (minimum 30 targets). The fraction recovered in the top 10 is, at *n* = 30, 0.38 for FSR and 0 for ARACNe3-top100; at *n* = 100, 0.84 and 0.2; at *n* = 400, 0.82 and 0.74 (ceiling with the true network, 0.86). ARACNe3 produces no edges at *n* = 30 and, on average, 14.6 edges at *n* = 60 with strong signal (none with weak signal), too few to evaluate. This agrees with the recommendation of at least 100 expression profiles for ARACNe, an empirical lower bound below which its accuracy drops rapidly (Margolin et al. [14]; see also [13, 3]). Implementation [1]; algorithm in Margolin et al. [13], Lachmann et al. [8] and Griffin et al. [6].

### Synthetic and real results at *n* ≈ 100

The two measurements look discordant at *n* ≈ 100: in synthetic data ARACNe3-top100 recovers few master regulators (0.2), while in real cohorts with *n* ≥ 95 it overlaps with FSR in 0.7 of the top 20. Two features of the design can explain part of the gap. The synthetic generators are linear–Gaussian in construction (the same regime that favours correlation in Section 5.2), and ARACNe3 builds each subnetwork on 63 % of the samples, so at *n* = 100 every subnetwork rests on about 63 tumours, below its recommended size. The real-data overlap, in turn, measures consistency between the two methods and not correctness of either. We do not claim that the synthetic result predicts real-data accuracy; we claim only that both methods give consistent heads of the ranking where both can be run, and that only FSR runs below *n* ≈ 100.

## 7 Minimum cohort size

FSR yields regulons of full size at any *n*, so regulon size cannot indicate whether a cohort is large enough. We define the lower bound empirically, by the sampling variability of the master-regulator ranking relative to the variability between inference methods.

### Why a bound is needed

Fixed-size regulons are an operational advantage and a risk. For a regulator with no true targets among about 14 000 candidates, the 100th-largest null |*ρ*_*S*_| is approximately 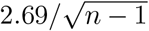 (Gaussian approximation to the null of *r*_*S*_, independent genes): 0.53 at *n* = 27, 0.35 at *n* = 60 and 0.27 at *n* = 100. The fixed threshold |*ρ*_*S*_| ≥ 0.2 does not control false positives at small *n*: the null probability of exceeding it is 0.31 at *n* = 27 and 0.12 at *n* = 60, and the threshold only becomes comparable to the top-100 cut at *n* ≈ 182. Below that size the truncation to 100 targets, not the threshold, is what limits the regulon, and the regulon of a regulator without signal is filled with chance associations. This is a heuristic argument that motivates an empirical bound; it is not a derivation of one.

### Design

5 large real cohorts (*n* from 190 to 562) were subsampled without replacement to *n ∈* { 20, 27, 40, 60, 80, 100, 150 }, 5 replicates each. An FSR network was inferred from each subsample and NaRnEA was run with the signature of the full cohort, which is held fixed so that only the network varies. The reference is the result with the network of the full cohort.

Because the signature is fixed, the relevant yardsticks are those in which the signature is also held fixed: the agreement between two inference methods on the same full cohort (FSR vs. ARACNe3-top100, 0.7), the agreement between the networks of two subtypes of the same cancer with the same signature (0.55), and the null (0.35).

### Results (Fig. 5)

Median top-20 overlap with the full-cohort result, with interquartile range over cohorts and replicates:

**Figure 5.**
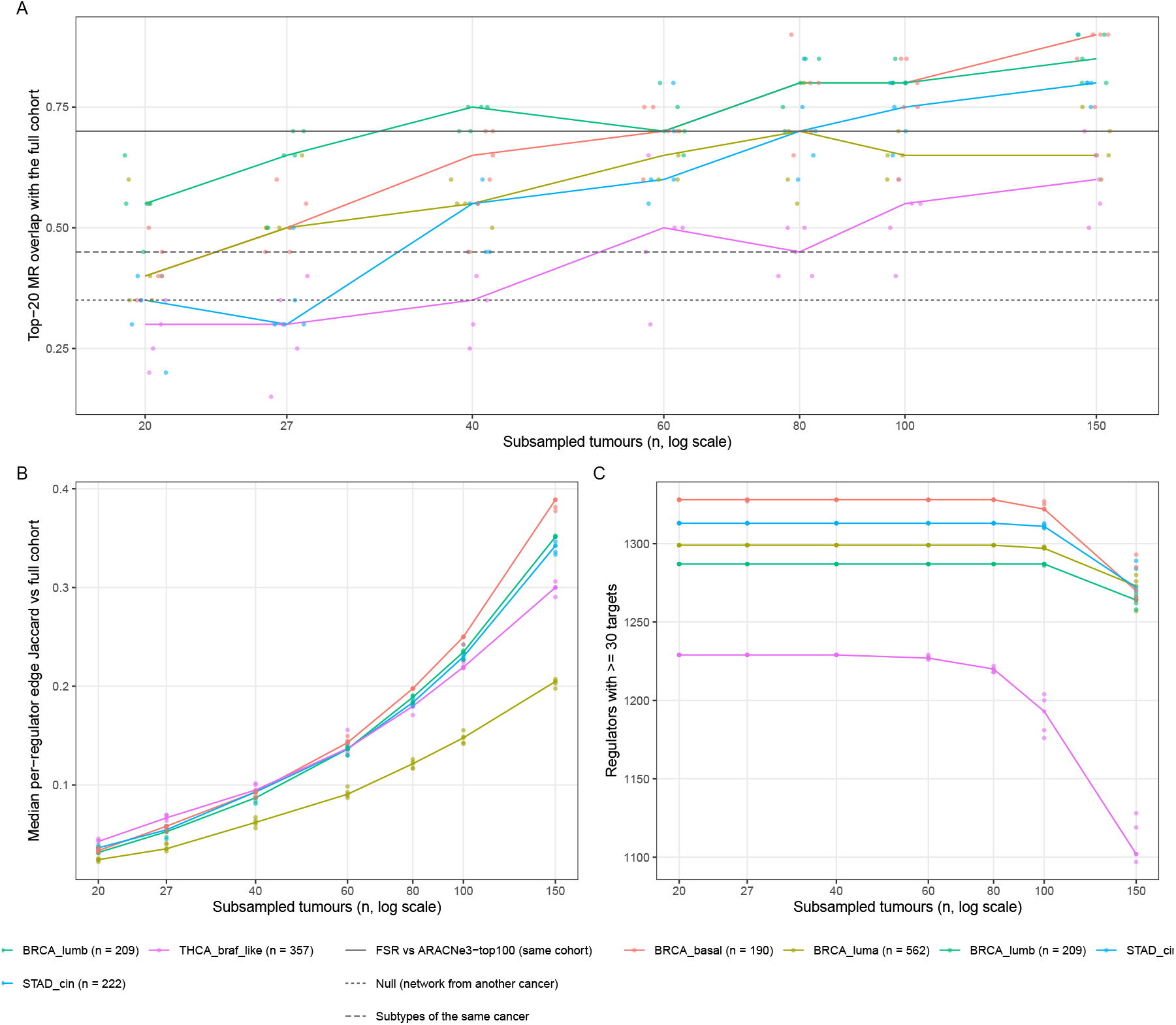
Subsampling of large real cohorts. (A) Top-20 overlap of master regulators with the full-cohort result as a function of *n*; horizontal lines mark the agreement between methods, between subtypes and under the null. (B) Per-regulator edge Jaccard index with the full-cohort network. (C) Regulators with enough targets for NaRnEA.

| *n* | 20 | 27 | 40 | 60 | 80 | 100 |
|---|---|---|---|---|---|---|
| median | 0.4 | 0.5 | 0.55 | 0.65 | 0.7 | 0.75 |
| IQR | 0.35–0.5 | 0.35–0.55 | 0.45–0.7 | 0.6–0.7 | 0.6–0.8 | 0.6–0.8 |

At *n* = 20–27 the median overlap of a network with the result of its own cohort falls below the agreement between the networks of two different subtypes, so such a network reproduces its own cohort’s master regulators less well than a network from another subtype reproduces a cohort’s. The method-to-method agreement is reached between *n* = 60 and *n* = 80; the two cannot be distinguished, because the overlap moves in steps of one regulator in twenty and their interquartile ranges overlap almost entirely. Edge-level stability is low at every *n* (median per-regulator Jaccard index with the reference 0.136 at *n* = 60), while the head of the ranking is stable: the ranking tolerates imprecise edges, as in the synthetic data. THCA BRAF-like, whose reference regulons are the smallest, is the least stable cohort (0.55 at *n* = 100).

### Robustness to *k*

To check that the conclusions do not depend on the choice of the top 20, each comparison was repeated for *k ∈* { 5, 10, 15, 20, 30, 40, 50}, with 95 % confidence intervals for the median obtained by bootstrap over the independent units (cohorts; cancers for pairs of subtypes of the same cancer, which share a cancer), 2 000 resamples. The curves for every comparison are in Supplementary Fig. S9. Agreement between methods stays at 0.6–0.7 over the whole range, the ordering methods *>* subtypes of the same cancer *>* null holds for 6 of 7 values of *k* and fails only for *k* ≤ 5, where a single regulator moves the overlap by 20 %, and the confidence interval of method agreement lies above that of between-subtype agreement for 6 of 7 values. The confidence intervals of between-subtype agreement and of the null overlap intersect at every *k*: with ten cancers as resampling units, that particular difference is not established. The smallest subsample size at which the median overlap reaches method agreement ranges from 60 to 100 across *k*.

### Operational rule

We take *n* ≥ 60 as the lower bound for a subtype-specific network: it is the smallest size at which sampling variability is of the order of the variability between methods. Because the criterion moves between 60 and 100 with *k*, we treat 60 ≤ *n <* 100 as a grey zone, in which results are usable but their sampling variability is still above that between methods, and *n* ≥ 100 as the range in which an ARACNe3 comparison is also feasible. The criterion is relative to the method yardstick, not an absolute accuracy, and four caveats apply: it rests on 5 cohorts; the signature is held fixed, so the additional noise of a signature estimated from few tumours is not included; the reference is itself an estimate; and subsamples of the smaller reference cohorts are nested in the reference to a degree that grows with *n* (at *n* = 150 a cohort of 190 tumours contributes most of its samples to both the subsample and the reference), which inflates the overlap at large *n*. The criterion supports excluding *n <* 60 more firmly than it singles out 60 over 80.

### Below the bound

A cohort under the bound could instead be interrogated with a network inferred from all tumours of its cancer. We tested this on 11 large cohorts reduced to 30 tumours (3 replicates), comparing three networks against the full-cohort reference (Fig. 6): the cohort’s own network from the 30 tumours, a cancer-wide network (the other subtypes plus the same 30 tumours), and the network of another subtype. The acceptance criterion, fixed in advance, was that the cancer-wide network outperform the own network (paired Wilcoxon test over cohorts, *p <* 0.05) and the network of another subtype. Median top-20 overlap was 0.6 (cancer-wide), 0.55 (own) and 0.433 (another subtype). The cancer-wide network outperformed the own network in 7 of 11 cohorts (*p* = 0.0502) and another subtype’s network in 10 (*p* = 0.00667), so it does not meet the criterion, although narrowly. With 11 cohorts the test has little power, so the result is the absence of evidence of a rescue, not evidence of its absence. The own network, in turn, outperforms the network of another subtype (0.55 vs. 0.433), so at *n* ≈ 30 it still carries subtype-specific information, even though its reliability is low. The cancer-wide network also homogenises subtypes: in lung squamous carcinoma, where three of four subtypes fall below the bound, the top-20 overlap between subtypes rises from 0.25–0.6 with their own networks to 0.7–0.85 with the cancer-wide network. This could reflect lost subtype specificity or shared biology that the subtype-specific networks fail to recover; the data do not separate the two. Cohorts below the bound are therefore analysed with their own networks and reported as exploratory.

**Figure 6.**
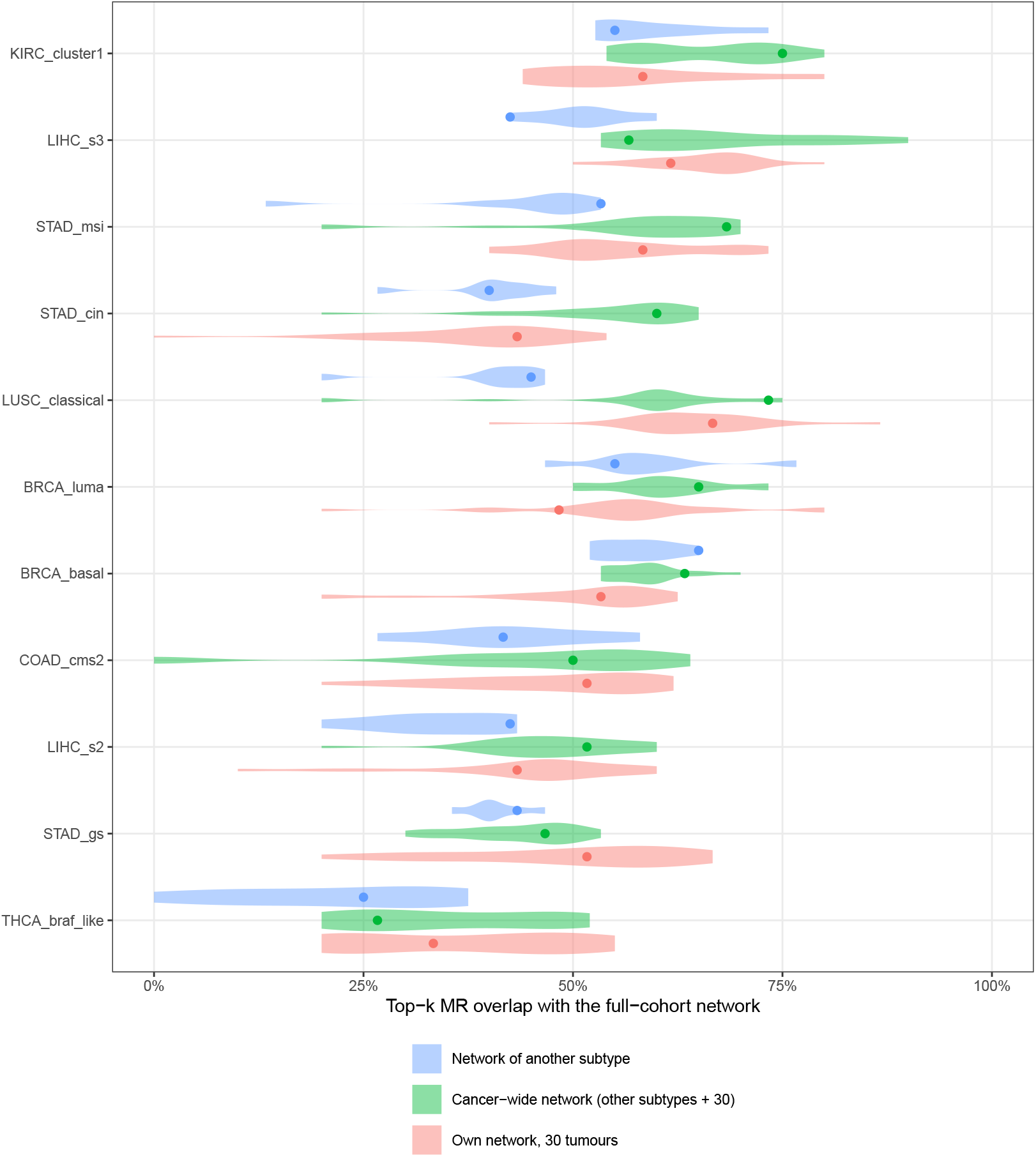
Large cohorts reduced to 30 tumours: overlap of the top-*k* master regulators with the full-cohort result for the cohort’s own network, a cancer-wide network and the network of another subtype of the same cancer. Violins: distribution over *k ∈* {5, 10, 15, 20, 30, 40, 50} and the 3 replicates (the network of another subtype has no replicates); points: *k* = 20, mean over replicates, the value quoted in the text.

**8 Discussion**

### What was conceded and what was gained

A homogeneous design built on rank correlation concedes three things, each quantified above: sensitivity to non-monotone dependence (at most about 0.05–0.10 of the recovered master regulators in a synthetic worst case, with unknown real prevalence), a heuristic per-target DPI whose decisions differ measurably from the triangle DPI and that is less conservative than ARACNe3 at *n* = 100, and a measurable method effect in the tail of the ranking. It gains a network at every cohort size, higher power than the MI estimators simulated at *n ≲* 200, and, above all, comparability across subtypes and cancers, the quantity the analysis is built to measure.

### Status of the claims

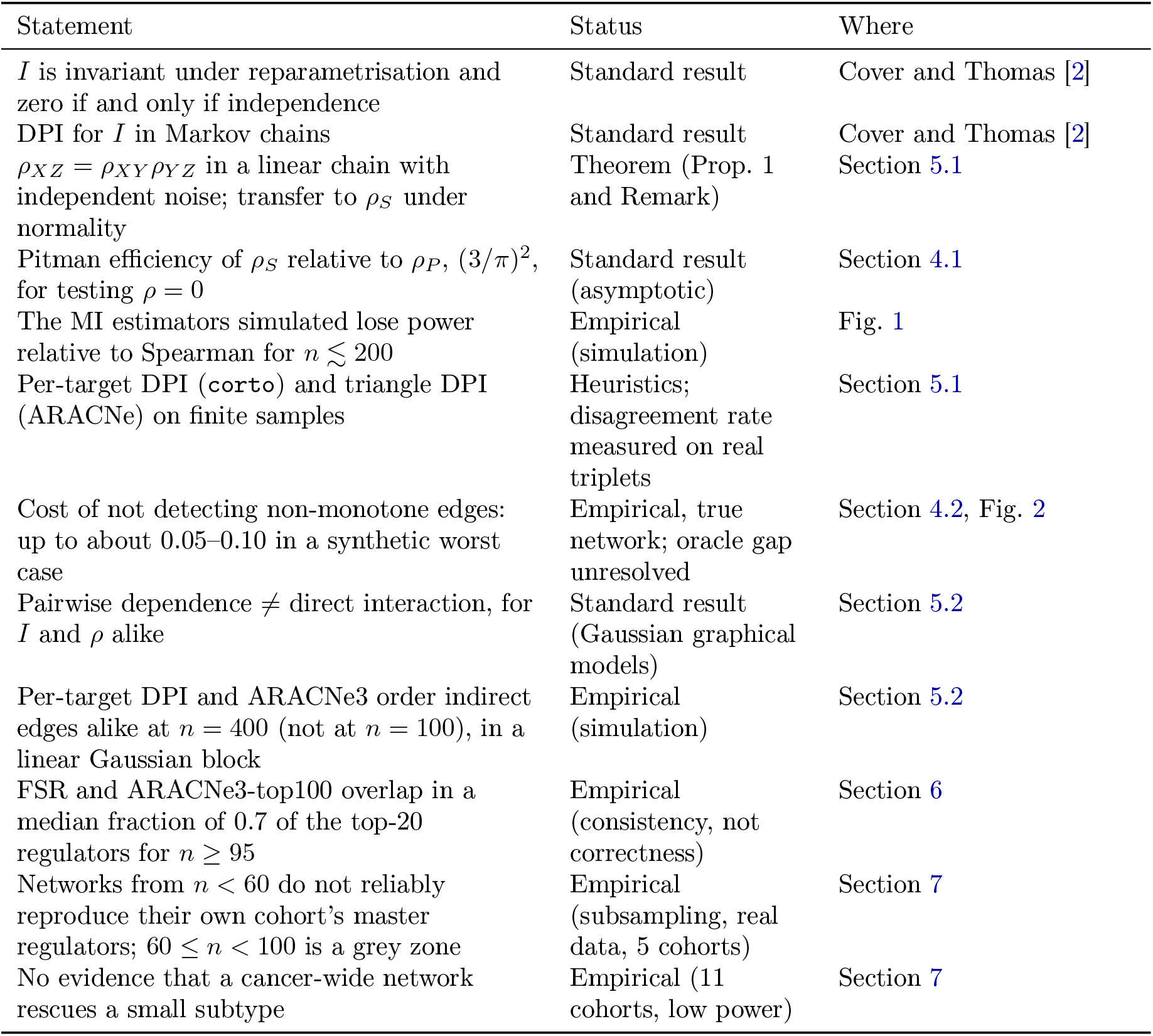

| Statement | Status | Where |
|---|---|---|
| *I* is invariant under reparametrisation and zero if and only if independence | Standard result | Cover and Thomas [2] |
| DPI for *I* in Markov chains | Standard result | Cover and Thomas [2] |
| ρ<sub>XZ</sub> = ρ<sub>XY</sub>ρ<sub>YZ</sub> in a linear chain with independent noise; transfer to ρ<sub>S</sub> under normality | Theorem (Prop. 1 and Remark) | Section 5.1 |
| Pitman efficiency of ρ<sub>S</sub> relative to ρ<sub>P</sub>, (3/π)<sup>2</sup>, for testing ρ = 0 | Standard result (asymptotic) | Section 4.1 |
| The MI estimators simulated lose power relative to Spearman for *n* ≲ 200 | Empirical (simulation) | Fig. 1 |
| Per-target DPI (corto) and triangle DPI (ARACNe) on finite samples | Heuristics; disagreement rate measured on real triplets | Section 5.1 |
| Cost of not detecting non-monotone edges: up to about 0.05–0.10 in a synthetic worst case | Empirical, true network; oracle gap unresolved | Section 4.2, Fig. 2 |
| Pairwise dependence ≠ direct interaction, for *I* and ρ alike | Standard result (Gaussian graphical models) | Section 5.2 |
| Per-target DPI and ARACNe3 order indirect edges alike at *n* = 400 (not at *n* = 100), in a linear Gaussian block | Empirical (simulation) | Section 5.2 |
| FSR and ARACNe3-top100 overlap in a median fraction of 0.7 of the top-20 regulators for *n* ≥ 95 | Empirical (consistency, not correctness) | Section 6 |
| Networks from *n* < 60 do not reliably reproduce their own cohort’s master regulators; 60 ≤ *n* < 100 is a grey zone | Empirical (subsampling, real data, 5 cohorts) | Section 7 |
| No evidence that a cancer-wide network rescues a small subtype | Empirical (11 cohorts, low power) | Section 7 |

#### Recommended use

Cohorts with *n* ≥ 100 can be analysed with FSR, and ARACNe3 can serve as a sensitivity check. Cohorts with 60 ≤ *n <* 100 are analysed with FSR and their rankings are interpreted with the sampling variability in mind. Cohorts with *n <* 60 are analysed with their own FSR networks and reported as exploratory.

### Limitations

1. FSR networks are pruned co-expression networks, not networks of direct interactions; so are ARACNe3 networks. Master regulators are interpreted as regulators whose *co-expressed* targets change coherently, and co-expressed TF families are interpreted at the family level. Confounders such as tumour purity and proliferation are not resolved by either method.
2. The per-target DPI of `corto` has no theoretical support beyond the linear case. In real chains without a direct edge it retains the indirect regulator in a fraction 0.182 of cases, and by construction it discards independent co-regulators of the same target, which limits recall. The triangle rule of `corto` ≥ 1.4.0 keeps them, but did not recover more master regulators in our simulations (Section 5.1).
3. Non-monotone dependencies exist that correlation does not detect, and their prevalence in these cohorts is unknown. Their small effect on the ranking depends on the current scoring method; it has been quantified and is not zero.
4. No method performs well with small cohorts. FSR produces full-size regulons at any *n*, but below *n* ≈ 60 they do not reproduce the master regulators of their own cohort reliably (Section 7); such cohorts are reported as exploratory.
5. The synthetic benchmarks are linear and Gaussian by construction and favour correlation-based methods; the real-data comparison with ARACNe3 measures consistency, not correctness.
6. The threshold *r*_min_ = 0.2 was selected after inspecting the results, and a grid of values is reported in the Supplementary Note. In most real cohorts the threshold is inactive; the procedure rests on the DPI pruning and on the truncation to 100 targets.

## Reproducibility

Figures and numbers in this work are produced by `scripts/network_benchmark/10_appendix_ statistics.R` and `11_appendix_numbers_tex.R` in the `FSR_Networks` repository; those quoted from the Supplementary Note (Sections 3 and 6) come from `network_benchmark_key_numbers. tsv`. Every number in the text is a macro in `numbers.tex`, generated by these scripts, except the closed-form null quantities of Section 7, defined at the top of this file.

## Use of AI tools

Claude Opus 5.5 (Anthropic) was used as a coding assistant and OpenAI Codex was also used for language editing. The authors are responsible for the scientific content and its verification.

